# Loss of postsynaptic NMDARs drives nanoscale reorganization of Munc13-1 and PSD-95

**DOI:** 10.1101/2024.01.12.574705

**Authors:** Poorna A. Dharmasri, Emily M. DeMarco, Michael C. Anderson, Aaron D. Levy, Thomas A. Blanpied

**Author notes:** **Corresponding Author:** Thomas A. Blanpied. **Author contributions:** P.A.D. and T.A.B. designed research; P.A.D., E.M.D., and A.D.L. performed research; P.A.D., E.M.D., and M.C.A. analyzed data; P.A.D. and T.A.B. wrote the paper.

## Abstract

Nanoscale protein organization within the active zone (AZ) and post-synaptic density (PSD) influences synaptic transmission. Nanoclusters of presynaptic Munc13-1 are associated with readily releasable pool size and neurotransmitter vesicle priming, while postsynaptic PSD-95 nanoclusters coordinate glutamate receptors across from release sites to control their opening probability. Nanocluster number, size, and protein density vary between synapse types and with development and plasticity, supporting a wide range of functional states at the synapse. Whether or how the receptors themselves control this critical architecture remains unclear. One prominent PSD molecular complex is the NMDA receptor (NMDAR). NMDARs coordinate several modes of signaling within synapses, giving them the potential to influence synaptic organization through direct protein interactions or through signaling. We found that loss of NMDARs results in larger synapses that contain smaller, denser, and more numerous PSD-95 nanoclusters. Intriguingly, NMDAR loss also generates retrograde reorganization of the active zone, resulting in denser, more numerous Munc13-1 nanoclusters, more of which are aligned with PSD-95 nanoclusters. Together, these changes to synaptic nanostructure predict stronger AMPA receptor-mediated transmission in the absence of NMDARs. Notably, while prolonged antagonism of NMDAR activity increases Munc13-1 density within nanoclusters, it does not fully recapitulate these trans-synaptic effects. Thus, our results confirm that NMDARs play an important role in maintaining pre- and postsynaptic nanostructure and suggest that both decreased NMDAR expression and suppressed NMDAR activity may exert distinct effects on synaptic function, yet through unique architectural mechanisms.

**Significance Statement:** Synaptic transmission is shaped by the trans-synaptic coordination of molecular ensembles required for neurotransmitter release and receptor retention, but how receptors themselves influence this critical architecture remains unclear. Using state-of-the-art super-resolution microscopy, we report that loss of NMDA receptors from excitatory synapses alters both pre- and postsynaptic nano-organizational features. Notably, pharmacological antagonism of NMDA receptors also alters presynaptic features, but without fully mimicking effects of the knockout. This suggests that both NMDA receptor activity and presence at the synapse exert retrograde influence on active zone organization. Because numerous disease and activity states decrease expression or function of NMDA receptors, our results suggest that distinct nanostructural states contribute to the unique functional status of synapses in these disorders.

## Introduction

Several properties of synaptic transmission depend on the precise, subsynaptic organization of key effector molecules (Gou et al., 2022). In the presynaptic active zone (AZ), the number of nanoclusters (NCs) of the protein Munc13-1 predicts the number of release sites within a synapse (Karlocai et al., 2021; Sakamoto et al., 2018). The molecular organization within these NCs and the organization of proteins around them is thought to control many aspects of release dynamics, such as vesicle priming and release probability (Aldahabi et al., 2022). Key proteins within the postsynaptic density (PSD) also form NCs, including glutamate receptors (Goncalves et al., 2020; Kellermayer et al., 2018; MacGillavry et al., 2013; Nair et al., 2013; Tang et al., 2016) and the primary excitatory synapse scaffold protein PSD-95 (Broadhead et al., 2016; Fukata et al., 2013; MacGillavry et al., 2013; Tang et al., 2016). PSD-95 NCs align receptors with release sites to strengthen synaptic transmission (Ramsey et al., 2021; Tang et al., 2016). Synaptic nanostructure is quite dynamic, capable of change on the order of minutes (MacGillavry et al., 2013; Nair et al., 2013; Ramsey et al., 2021; Tang et al., 2016), and taking on different characteristics depending on cell identity (Dharmasri et al., 2023) and throughout development (Sun et al., 2022). Notably, nanostructural changes can support a wide range of unique functional states in the synapse (Chen et al., 2018; Glebov et al., 2016; Han et al., 2022). Despite the importance of synaptic nanostructure, we still have an incomplete understanding of the factors that influence subsynaptic organization.

NMDA receptors (NMDARs) are well positioned to act as modifiers of synaptic nanostructure, either as signaling molecules or structural organizers. NMDARs are present at nearly all synapses and their activity is involved in synaptogenesis (Hansen et al., 2021). NMDAR signaling has already been implicated in nanostructural rearrangements, as NMDAR-dependent LTP enlarges PSD-95 and AMPA receptor nanoclusters (Clavet-Fournier et al., 2023) and enhances the enrichment of PSD-95 within NCs as well as across from release sites (Tang et al., 2016). Additionally, LTD triggered by NMDA application conversely reduces PSD-95 NC number and alignment with release sites (Tang et al., 2016). Beyond receptor activity, NMDARs may also play a structural role within the synapse. Non-ionotropic actions of NMDARs impact spine stability (Alvarez et al., 2007) and gate synaptic plasticity (Dore et al., 2017). Furthermore, NMDARs exist in supercomplexes that are associated with multiple scaffold molecules (Frank et al., 2016). Their intracellular C-termini coordinate multiple PSD-95 molecules (Zeng et al., 2018) and enhance the ability of other molecules to phase separate (Yang et al., 2023), possibly allowing them to directly alter PSD nanostructure. In addition, NMDARs interact with EphB2 (Hanamura et al., 2017; Washburn et al., 2020) and Nlgn1 (Budreck et al., 2013) in the synaptic cleft, giving them the potential to act as, or to modify, structural linkers between the pre- and postsynapse and thereby influence active zone organization.

Here we tested whether NMDARs influence synaptic nanostructure by measuring how NMDAR loss influences the nanoscale organization of Munc13-1 and PSD-95. Our findings indicate that loss of NMDARs yields larger synapses with substantially altered nanostructure, including an increase in NC number and protein density within NCs on both sides of the synapse. More nuanced changes were also present, including an increase in the number of release sites aligned with PSD-95 and an alteration in the relationship between active zone size and release site number. Surprisingly, the trans-synaptic effects of NMDAR loss were not fully recapitulated by a loss of NMDAR activity. Our data suggest that the NMDAR complex regulates trans-synaptic nanostructure by both structural and signaling mechanisms.

## Materials and Methods

### DNA constructs

LentiCRISPRv2GFP (LCv2) was a gift from David Feldser (Addgene plasmid #82416; http://n2t.net/addgene:82416; RRID:Addgene_82416) and is a lentiviral vector that contains a gRNA scaffold and a downstream Cas9-P2A-GFP cassette to enable gRNA, Cas9, and cell identification marker expression from a single virus. gRNA against *GRIN1* was cloned into LCv2 using a protocol from the Zhang lab (https://media.addgene.org/cms/filer_public/4f/ab/4fabc269-56e2-4ba5-92bd-09dc89c1e862/zhang_lenticrisprv2_and_lentiguide_oligo_cloning_protocol_1.pdf). Primers for the gRNA against *GRIN1* were designed as previously described (*GRIN1*#2 from Incontro et al., 2014): Forward (5’ to 3’) CACCGACTAGGATAGCGTAGACCTG; Reverse (5’ to 3’) AAACCAGGTCTACGCTATCCTAGTC. psPAX2 (Addgene plasmid #12260; http://n2t.net/addgene:12260; RRID:Addgene_12260) and pMD2.G (Addgene plasmid #12259; http://n2t.net/addgene:12259 RRID:Addgene_12259) were gifts from Didier Trono. pFCaGW was generated as described in (Dharmasri et al., 2023).

### Lentivirus production

Lentiviruses of LCv2 containing either No Guide or gRNA against *GRIN1* were generated using HEK293T cells (ATCC CRL-3216), essentially as in (Dharmasri et al., 2023). Briefly, HEK293T cells were transfected with 6 μg of either lentiviral construct, 4 ug psPAX2, and 2 μg pMD2.G using PEI, and 4 - 6h later the media was exchanged for neuron culture media. Virus-conditioned media was harvested after 48h, centrifuged briefly and 0.45 μm PES-filtered to remove debris, and stored at -80°C in single-use aliquots.

### Primary neuron culture

All animal procedures were approved by the University of Maryland Animal Use and Care committee. Dissociated hippocampal neuron cultures were prepared from E18 Sprague-Dawley rat embryos (Charles River) of both sexes as described (Dharmasri et al., 2023). Neurons were plated on poly-L-lysine-coated coverslips at 30,000 cells/coverslip (18 mm #1.5, Warner Instruments) in Neurobasal A + GlutaMax, gentamycin, B27 supplement, and 5% FBS. After 24 hours, media was changed to the same but lacking FBS, and after 1 week supplemented with an additional half volume of media + FUDR to suppress glial growth. For knockout experiments, neurons were infected at DIV 1 with 250 μl of unconcentrated No Guide or *GRIN1*-targeting LCv2 lentivirus and fixed at DIV22 for both confocal and DNA-PAINT experiments. For APV experiments, neurons were infected at plating as in (Metzbower et al., 2019) with pFCaGW to express GFP in a subset of neurons.

### Drug Treatments

D,L-APV (Sigma) dissolved in dH_2_O and stored at 20 mM was diluted 1:200 into the culture media for a final concentration of 100 μM. An equal volume (7.5 μL/well) of dH_2_O was added to parallel coverslips for vehicle control. Treatment was started on DIV 14 and additional dosages of D,L-APV and vehicle were added to coverslips every 48 hours through DIV 20 prior to fixation on DIV 21.

### Antibody-dye conjugation

Secondary antibody for confocal imaging of PSD-95 was assembled in house as in (Dharmasri et al., 2023). Briefly, donkey anti-mouse IgG2a was mixed with NHS-Cy3B (GE PA63101) at ∼13:1 molar ratio of dye:IgG for 1h at RT to achieve a final dye/IgG ratio of ∼3:1, and excess dye was removed by Zeba desalting column (Thermo). Conjugated antibody was diluted to ∼1.25 mg/mL in 50% glycerol, aliquoted, and stored at -20°C.

### Immunostaining

LCv2 No Guide and LCv2 *GRIN1* lentivirus-infected coverslips or vehicle and APV treated coverslips from the same plate were fixed with 4% PFA + 4% sucrose in PBS for 8 minutes at room temperature (RT), washed 3 x 5 minutes with PBS + 100 mM glycine (PBSG), permeabilized 20 minutes RT with 0.3% Triton X-100 in PBSG, and blocked 20 minutes RT with 10% donkey serum + 0.2% Triton X-100 in PBSG.

For confocal imaging, the neurons were stained overnight at 4°C with primary antibodies mouse IgG2A anti-PSD-95 and either rabbit anti-Munc13-1 or rabbit anti-GluN1 in 10% donkey serum + 0.2% Triton X-100 in PBSG. The next day, cells were washed 3 x 5 minutes in PBSG, then incubated with the secondary antibodies goat anti-mouse IgG2a Cy3B, donkey anti-rabbit AlexaFluor647, and GFP-Booster nanobody for 1 hour at RT in PBSG. Finally, the cells were washed 3 x 5 minutes in PBSG, post-fixed with 4% PFA + 4% sucrose in PBS for 20 minutes and washed 3 x 5 minutes in PBSG again before storage at 4°C until imaging.

For DNA-PAINT, PSD-95 and Munc13-1 were stained with primaries preincubated (Sograte-Idrissi et al., 2020) with custom-made single-domain antibodies (sdAbs; Massive Photonics) carrying one of two oligonucleotide docking strands optimized for DNA-PAINT, as described (Anderson et al., 2023; Strauss and Jungmann, 2020). Briefly, the primary antibodies against PSD-95 and Munc13-1 were incubated separately with a 2.5-fold molar excess of anti-mouse sdAb-F1 or anti-rabbit sdAb-F3, respectively, for 20 minutes at RT, to saturate the antibody with sdAb. Rabbit Fc fragment was added to the Munc13-1 incubation at 2-fold molar excess for a further 20 minutes to remove unbound nanobody. Both preincubations were then diluted to their final working concentrations in 10% donkey serum + 0.2% Triton X-100 in PBSG and incubated on the cells overnight at 4°C. The next day, the cells were processed as for confocal imaging, but including only the GFP-Booster. 90 nm gold nanoparticles (Cytodiagnostics) were added at 1:10 dilution for 10 minutes before imaging as fiducials for drift and chromatic aberration correction.

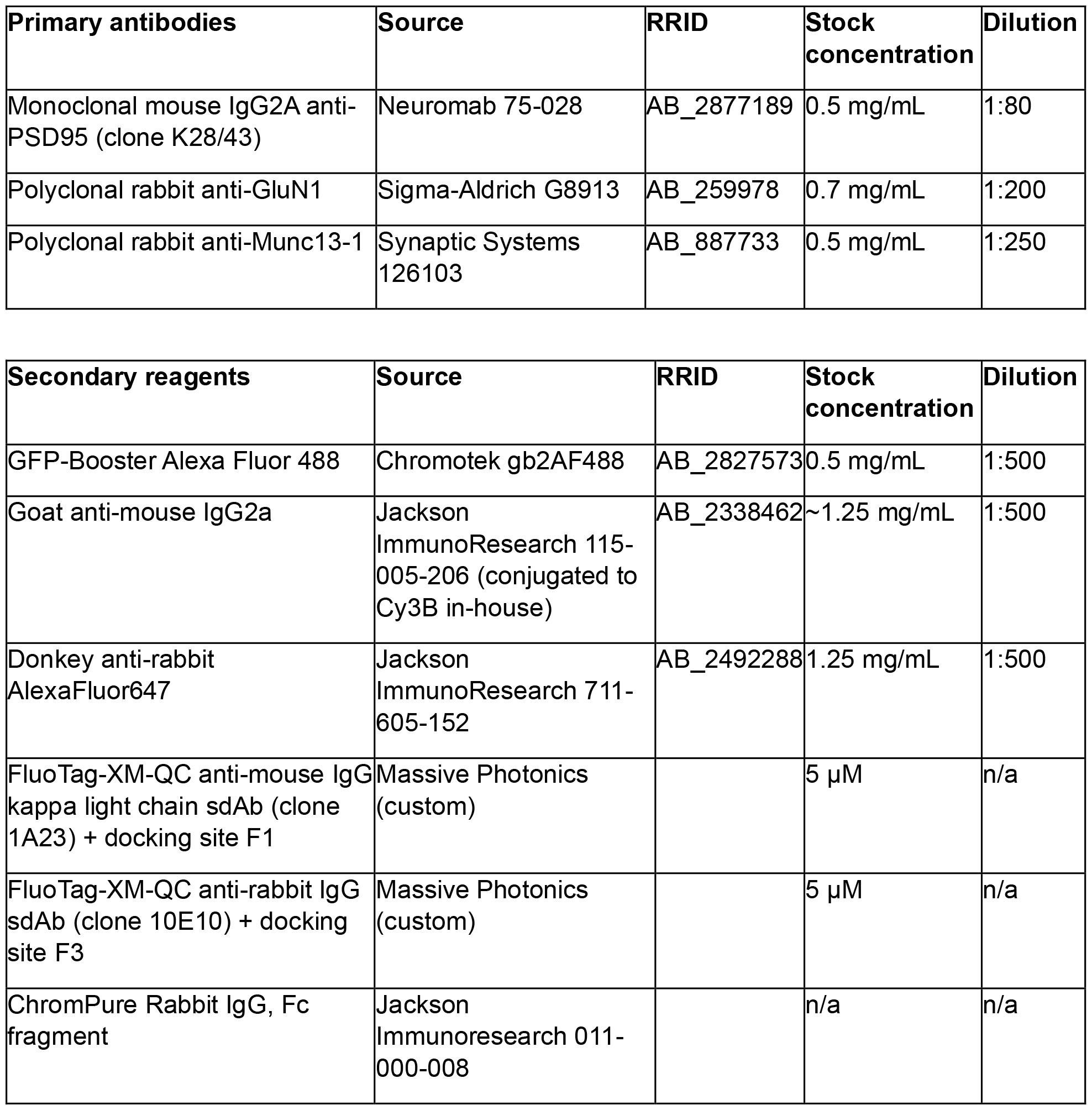

| Primary antibodies | Source | RRID | Stock concentration | Dilution |
| --- | --- | --- | --- | --- |
| Monoclonal mouse IgG2A anti-PSD95 (clone K28/43) | Neuromab 75-028 | AB_2877189 | 0.5 mg/mL | 1:80 |
| Polyclonal rabbit anti-GluN1 | Sigma-Aldrich G8913 | AB_259978 | 0.7 mg/mL | 1:200 |
| Polyclonal rabbit anti-Munc13-1 | Synaptic Systems 126103 | AB_887733 | 0.5 mg/mL | 1:250 |

| Secondary reagents | Source | RRID | Stock concentration | Dilution |
| --- | --- | --- | --- | --- |
| GFP-Booster Alexa Fluor 488 | Chromotek gb2AF488 | AB_2827573 | 0.5 mg/mL | 1:500 |
| Goat anti-mouse IgG2a | Jackson ImmunoResearch 115-005-206 (conjugated to Cy3B in-house) | AB_2338462 | ~1.25 mg/mL | 1:500 |
| Donkey anti-rabbit AlexaFluor647 | Jackson ImmunoResearch 711-605-152 | AB_2492288 | 1.25 mg/mL | 1:500 |
| FluoTag-XM-QC anti-mouse IgG kappa light chain sdAb (clone 1A23) + docking site F1 | Massive Photonics (custom) | | 5 μM | n/a |
| FluoTag-XM-QC anti-rabbit IgG sdAb (clone 10E10) + docking site F3 | Massive Photonics (custom) | | 5 μM | n/a |
| ChromPure Rabbit IgG, Fc fragment | Jackson Immunoresearch 011-000-008 |  | n/a | n/a |

### Confocal microscopy

Confocal images in knockout experiments were acquired on an Olympus IX81 inverted microscope with a 60x/1.42 NA oil immersion objective. Confocal images in APV experiments were acquired on a Nikon TI2 Eclipse inverted microscope with a Nikon Apo TIRF 60x/1.49 NA objective. All images were acquired using a Dragonfly confocal unit (Andor). Excitation laser light (488, 561, or 638 nm) from an Andor ILE, flattened by an Andor Beam Conditioning Unit, was passed to the sample by a 405/488/561/640 quadband polychroic (Chroma). Emission light was passed through an appropriate bandpass filter (ET525/50, ET600/50 (Chroma), or Em01-R442/647 (Semrock), for AlexaFluor488, Cy3B, and AlexaFluor647 emission, respectively) and collected on a Zyla 4.2 sCMOS camera (Andor). Cells of interest were imaged with confocal z-stacks with 0.5 μm z-steps at 50-80% laser power (∼1-2 W/cm^2^) with 200 ms exposure, with each channel imaged sequentially.

### Confocal analysis

Confocal images were analyzed using a custom FIJI (Schindelin et al., 2012) macro. For each analysis at least 4 cells from one to three independent culture weeks were analyzed. One region of dendrite containing at least 20 synapses was cropped from each cell. A binary mask was then applied using the GFP signal to isolate synapses from cells of interest (e.g. expressing LCv2). To isolate ROIs from individual synapses within the GFP binary, two masks were applied using the PSD-95 and Munc13-1 signal. All thresholds were user-defined due to differences in puncta intensity. Only PSD-95 and Munc13-1 ROIs that had overlap were then used for synaptic measurements. Mean and integrated intensity were then calculated, background subtracted and normalized to the associated control condition for each culture week. Puncta area was also calculated using the imaged pixel size of 103 nm. For the GluN1 KO validation experiments, PSD-95 ROI masks within the GFP binary were used to isolate ROIs and measure GluN1 signal.

### Single-molecule microscopy

3D DNA-PAINT images in the knockout experiments were acquired on a custom microscope built around an RM21 base (Mad City Labs) that has been previously described (Dharmasri et al., 2023).

3D DNA-PAINT images in the APV experiments were acquired on a Nikon TI2 inverted microscope with a 100x/1.49 NA Apo TIRF oil immersion objective. Excitation light was supplied by LBX-488-100 (488 nm; Oxxius), SAP-561-300 (561 nm; Coherent), and LCX-640-300 (640 nm; Oxxius) lasers managed by an L6Cc laser combiner (Oxxius). The combiner was fiber-coupled to a manual Nikon TIRF illuminator used to adjust the incidence angle to achieve Highly Inclined and Laminated Optical (HILO) illumination and reflected to the sample via a ZT405/488/561/640/rpcv2-UF2 quadband polychroic (Chroma). Emission light was passed through a MicAO adaptive optics device (Imagine Optic) to correct aberrations in the point-spread function and introduce astigmatism, followed by a DV2 image splitter (Photometrics) equipped with a T640lpxr dichroic and ET655lp single band and 59044m dual band emission filters (Chroma). Emission was collected on an Ixon+ 897 EM-CCD camera (Andor). Z-stability was maintained by the Nikon Perfect Focus System. The microscope, lasers, and camera were controlled by Nikon Elements software, and the MicAO by separate Imagine Optic software.

### Single-molecule imaging

GFP cells were identified with low power 488 nm laser. F1-Atto643 (for PSD-95) and F3-Cy3B (for Munc13) imagers were diluted in imaging buffer (1x PBS pH7.4 + 500 mM NaCl + PCA/PCD/Trolox oxygen scavengers; Schnitzbauer et al., 2017) to 0.25 nM each (for knockout experiments and PSD-95 in APV experiments) or 0.125 nM (for Munc13-1 in APV experiments). Diluted imagers were added to the sample, which was allowed to equilibrate for at least 10 minutes to reduce drift. For knockout experiments, 40,000 frames were acquired with 150 ms exposure, with lasers power densities at the sample of 0.10 kW/cm^2^ for the 638 nm laser and 0.059 kW/cm^2^ for the 561 nm laser. The 785 nm laser was set to 50% power and used for focus lock. For APV experiments, 50,000 frames were acquired with 100 ms exposure, with laser power densities at the sample of 0.046 kW/cm2 for the 561 nm laser and 0.065 kW/cm2 for the 640 nm laser. Imaging buffer was refreshed between regions.

### Single-molecule localization

For the knockout experiments, molecule locations were determined using the Super-resolution Microscopy Analysis Platform, SMAP (Ries, 2020), identically to (Dharmasri et al., 2023). Briefly, 3D calibrations were generated from z-stacks of 100 nm TetraSpeck beads using the *calibrate3DsplinePSF* plugin. TetraSpeck beads imaged without astigmatism for 100 frames were localized using the *PSF free* fitter with ROI size 7 and used for dual-view chromatic aberration correction. For 3D experimental images, the *spline* fitter was used with ROI size 15, adjusting the RI mismatch to 0.83 as appropriate for the system, and loading the previously calculated 3D calibration.

For the APV experiments, molecule localizations were determined using Picasso (Schnitzbauer et al., 2017). Z-stacks of the 3D PSF of 100 nm TetraSpeck beads on coverglass were captured with 20 nm step size for each wavelength and the 3D calibration generated in Picasso Localize using the *Calibrate 3D* function. The z-stack was adjusted such that the middle frame represented 0 nm in relative z position and that the extremes of the z stack represented +/-400 nm. Images output from Elements were loaded into FIJI, cropped to separate each wavelength due to simultaneous imaging with the DV2, and converted to a .*raw* format prior to loading into Picasso Localize. Images were localized using the LQ, Gaussian method and GPUfit using a wavelength-specific 3D calibration and a magnification factor of 0.83, adjusting the net gradient on a per image basis to account for idiosyncrasies in background.

### Single-molecule analysis

All analysis was conducted using custom routines in MATLAB (Mathworks) which relied, in part, on command line calls to Picasso for specific functions.

#### Processing of super-resolution images

Super-resolution images were processed identically to (Dharmasri et al., 2023). Briefly, Drift was first corrected by Picasso Render’s *Undrift by RCC* function. Chromatic aberrations were corrected using a polynomial transformation with MATLAB’s *fitgeotrans* function, calculated from the 2D bead images. Any residual offset of channels was corrected by cross-correlation. Poorly fit molecules were eliminated if their: localization precision was greater than 20 nm, PSF standard deviation was greater than 2 pixels, photon count was smaller than the mode of the whole-field histogram, or, for data analyzed in SMAP, if relative log likelihood was less than the first shoulder in the histogram (∼-1.5). Localizations were merged temporally using Picasso Render’s *Link localizations* function. Finally, localizations artificially moved to an extreme ceiling or floor in z by SMAP (Dharmasri et al., 2023) were eliminated per image.

#### Identifying putative synapses

Synapses were automatically segmented as in (Dharmasri et al., 2023). Briefly, Picasso Render’s *Clustering>DBSCAN* function was used with radius of 30 nm and minimum density of 5 to identify objects, where were filtered on the mean and standard deviation of the frames in which localizations within each object were present to eliminate gold fiducials or transient blinking of individual imager strands. Putative synaptic clusters whose 2D projection lacked any localizations of the opposing protein were eliminated. PSD-95 and Munc13-1 localizations were then treated as if from the same population of molecules and segmented into putative synapses using MATLAB’s *dbscan* function with epsilon of 30 nm and minpts of 4 for knockout experiments and an epsilon of 48 nm and minpts of 40 for APV experiments. Putative synapses were inspected for proper segmentation, with manual segmentation used as necessary. In knockout experiments, synapses with fewer than 60 localizations, or whose long/short axis ratio was greater than 2.5, or whose area was less than 1.5 pixels squared or larger than 20 pixels squared were removed; in APV experiments, these filters were synapses containing fewer than 50 localizations in either channel, long/short axis ratio greater than 2.5, or area less than 1 pixel squared. The remaining putative synapses were then judged and sorted for cis- and/or trans-synaptic analyses quality based on sampling density, corresponding presence and shape between both Munc13-1 and PSD-95 clusters, and z spread of localizations. Putative outlier localizations were removed by calculating the mean x,y, and z position of each point cloud and keeping only localizations within 2 standard deviations on each axis.

#### Quantitative analysis of synaptic nanostructure

Autocorrelation, synaptic and nanocluster volume, percentage of synaptic volume occupied by NCs, auto-enrichment, enrichment index, and percentage NCs aligned analyses were conducted as described in (Dharmasri et al., 2023). NC detection was also conducted as in (Dharmasri et al., 2023), with pairwise distance between each localization computed and input into the *dbscan* function using the optional inputs (‘Distance’,’precomputed’). *Epsilon* for each point cloud was determined first by calculating the mean minimal distance for each point within the point cloud and multiplying by a protein species-specific, empirically determined factor (in this study, 1.6 for Munc13-1 and 2.4 for PSD-95). Local density heat maps shown in Figure 2, 3, and 5 were generated using the calculated *epsilon* for each synapse and thus represent the input to DBSCAN. *Minpts* was set as previously described.

### Statistical analysis

Statistical analysis was conducted using GraphPad PRISM. Data were tested for normality using D’Agostino & Pearson, Shapiro-Wilk, and Kolmogorov-Smirnov tests. Data that passed normality checks were tested using two-tailed t-tests if variance was equal between groups or Welch’s t-test if not. Data that did not pass these checks were tested with two-tailed Mann-Whitney tests. No statistical methods were used to predetermine sample size. Experimenters were blinded during analyses.

## Results

### Loss of NMDARs Leads to Synaptic Enlargement

To test the impact of NMDAR loss on synaptic structure, we used CRISPR/Cas9 to genetically delete the obligate NMDAR subunit GluN1, which is expected to result in total loss of all NMDAR subtypes. We infected dissociated rat hippocampal neurons at DIV 1 with lentivirus (LentiCRISPRv2 (LCv2); Sanjana et al., 2014) expressing either no gRNA (No Guide; NG) or a previously validated gRNA against *GRIN1* (N1KO; Incontro et al., 2014), along with spCas9, and EGFP as a cell selection marker (Walter et al., 2017). We first verified the GluN1 knockout in our hippocampal cultures using immunocytochemistry (Fig 1A-B). While most cells showed robust loss of GluN1 two weeks after infection, we observed a subpopulation that still had synaptic GluN1 signal (Fig 1C, top). 21.05% of infected cells contained synaptic GluN1 levels within one standard deviation of the normalized mean synaptic content of control cells. This subpopulation was largely eliminated three weeks after infection (Fig 1C, bottom), with only 2.94% of cells with synaptic GluN1 levels within the same threshold. We thus conducted all experiments at this time point, fixing neurons at DIV 22 and staining for both Munc13-1 and PSD-95 for either confocal or 3D DNA-PAINT imaging.

**Figure 1.**
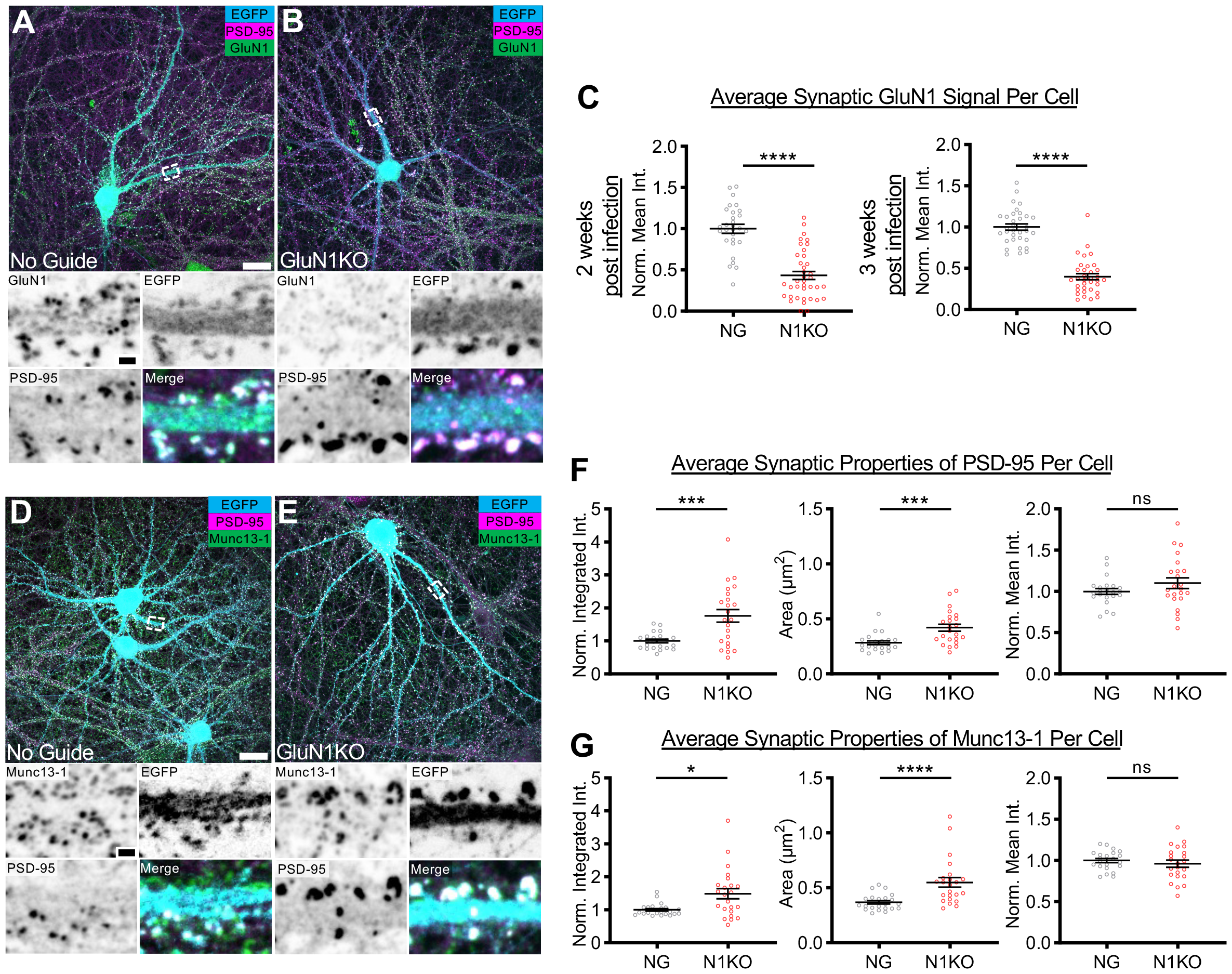
LCv2-*GRIN1* results in GluN1 loss and enlarged synapses three weeks post-infection. **A**,**B)** (Top) Example neurons in dissociated hippocampal cultures infected with either (A) LCv2-No Guide (NG) or (B) LCv2-*GRIN1* (N1KO). Scale bar: 20 μm. (Bottom) Dendrite stretches from example cells above showing GluN1 staining, PSD-95 staining, cell identifying marker, and merged image. Scale bar: 2 μm. **C)** Quantification of synaptic GluN1 mean intensity per imaged cell (NG, 1.00 +/-0.05, n = 29 cells; N1KO, 0.43 +/-0.05, n = 38 cells; p<0.0001 for 2 weeks post infection; NG, 1.00 +/-0.04, n = 33 cells; N1KO, 0.40 +/-0.04, n = 34 cells; p<0.0001 for 3 weeks post infection). **D**,**E)** (Top) Example neurons in dissociated hippocampal cultures infected with either (A) NG or (B) N1KO. Scale bar: 20 μm. (Bottom) Dendrite stretches from example cells above showing Munc13-1 staining, PSD-95 staining, cell identifying marker, and merged image. Scale bar: 2 μm. For F and G, per cell analyses were conducted to account for genotype (NG, n = 22 cells; N1KO, n = 23 cells). **F)** Quantification of synaptic PSD-95 properties per cell. (Left) Normalized integrated intensity (NG, 1.00 +/-0.05; N1KO, 1.76 +/-0.19; p=0.0007). (Middle) Puncta area (NG, 0.28 +/-0.02 μm^2^; N1KO, 0.42 +/-0.03 μm^2^; p=0.0003). (Right) Normalized mean intensity (NG, 1.00 +/-0.04; N1KO, 1.10 +/-0.06; p=0.3267). **G)** Quantification of synaptic Munc13-1 properties per cell. (Left) Normalized integrated intensity (NG, 1.00 +/-0.04; N1KO, 1.49 +/-0.15; p=0.0111). (Middle) Puncta area (NG, 0.37 +/-0.02 μm^2^; N1KO, 0.55+/-0.04 μm^2^; p<0.0001). (Right) Normalized mean intensity (NG, 1.00 +/-0.03; N1KO, 0.96 +/-0.04; p=0.4356). All data mean +/-SEM.

Visual inspection of GluN1 knockout neurons via confocal imaging revealed the presence of larger, brighter synapses than on control neurons (Fig 1D-E). To quantify whether NMDAR loss altered synapse properties, we analyzed a representative subpopulation of synapses from each imaged cell, normalizing to the control within each culture replicate. We found that GluN1 knockout cells contained PSDs with more PSD-95 than controls, as measured by integrated intensity (Fig 1F, left). This increase was due to an enlargement in PSD size (Fig 1F, middle) and not due to a change in protein density, as PSD-95 mean intensity was unchanged (Fig 1F, right). Intriguingly, we observed similar changes to presynaptic Munc13-1. AZs in synapses forming onto GluN1 knockout neurons contained more Munc13-1 (Fig 1G, left), which was again due to an enlarged AZ size (Fig 1G, middle) with no change to Munc13-1 density (Fig 1G, right). Together, this evidence suggests that NMDAR presence not only restricts PSD size (Ultanir et al., 2007), but AZ size as well.

### GluN1 Deletion Causes PSD-95 Nanostructure Remodeling

We next asked whether synaptic nanostructure was altered in these enlarged NMDAR-lacking synapses. We first analyzed the nanoscale organization of PSD-95 (Fig 2A). GluN1 knockout resulted in larger PSDs (Fig 2B), consistent with the larger puncta measured with confocal imaging. An autocorrelation analysis, which describes the heterogeneity of a protein’s subsynaptic distribution (Dharmasri et al., 2023; Ramsey et al., 2021; Tang et al., 2016), revealed a clear difference in PSD-95 organization when NMDARs were absent (Fig 2C). The higher magnitude over a larger shift radius is usually indicative of larger, denser PSD-95 NCs (Dharmasri et al., 2023). However, while an auto-enrichment analysis, which measures the average normalized density of a protein around its NC peak, does indicate an increase in local PSD-95 density (Fig 2D), we detected smaller PSD-95 NCs in GluN1 knockout neurons (Fig 2E). The autocorrelation analysis reflects multiple levels of organization within the synaptic protein cluster, whereas our NC detection method depends on a density threshold. Therefore the broadening of the curve could reflect an elevated density within the PSD that is below our NC detection threshold.

**Figure 2.**
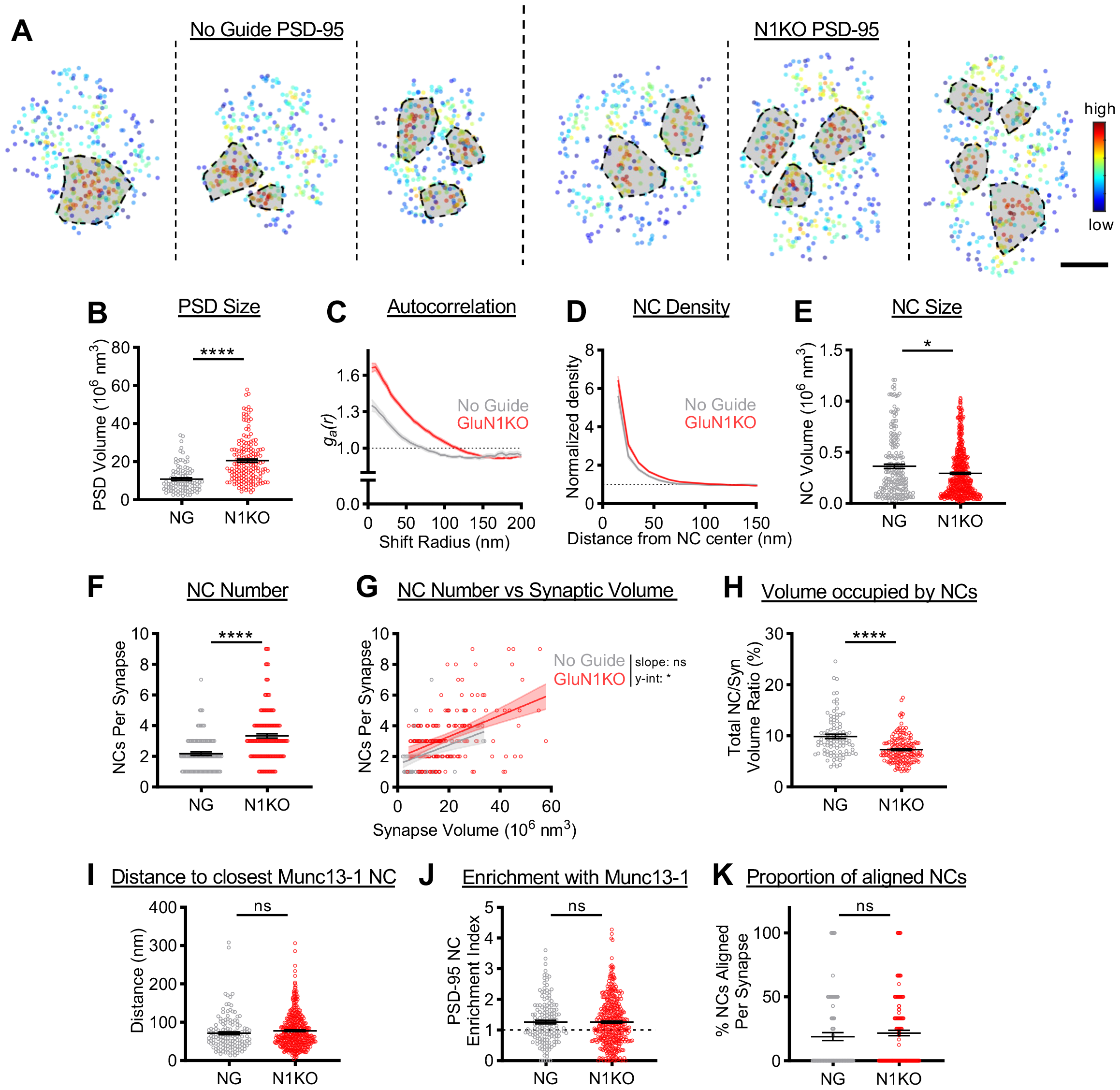
GluN1 knockout yields distinct PSD-95 nanoscale organization. **A)** Examples of 3D DNA-PAINT PSD-95 synaptic clusters from (Left) NG and (Right) N1KO synapses. Peak density-normalized local density indicated by heat maps. Detected NCs indicated by shaded shapes. Scale bar: 100 nm. Data in B,C,F-H are all synapses (NG, n = 96 synapses; N1KO, n = 169 synapses). Data in D,E,I-K are all NCs (NG, n = 196 NCs; N1KO, n = 511 NCs). **B)** PSDs are larger at N1KO synapses (NG, 10.83 +/-0.72 10^6^ nm^3^; N1KO, 20.54 +/-0.90 10^6^ nm^3^; p<0.0001). **C)** Autocorrelation predicts PSD-95 forms larger, denser NCs in N1KO synapses. **D)** Average PSD-95 NC has increased local density in N1KO synapses. **E)** PSD-95 NC volume is unexpectedly smaller with N1KO (NG, 0.36 +/-0.02 10^6^ nm^3^; N1KO, 0.29 +/-0.01 10^6^ nm^3^; p=0.0318). **F)** N1KO PSDs contain more PSD-95 NCs (NG, 2.18 +/-0.12 NCs/synapse; N1KO, 3.34 +/-0.14 NCs/synapse; p<0.0001). **G)** N1KO synapses have more PSD-95 NCs when accounting for PSD size (NG, slope: 0.063, y-intercept: 1.49, R^2^: 0.15; N1KO, slope: 0.069, y-intercept: 1.92, R^2^: 0.20. slope p=0.8011, y-intercept p=0.0134). **H)** N1KO results in less synaptic volume occupied by PSD-95 NCs (NG, 9.90 +/-0.41% volume; N1KO, 7.32 +/-0.20% volume; p<0.0001). **I)** PSD-95 NC peak’s distance to Munc13-1 NC peak is unchanged with N1KO (NG, 71.52 +/-3.78 nm; N1KO, 77.93 +/-2.23 nm; p=0.0876). **J)** Enrichment of Munc13-1 across from PSD-95 NCs is unchanged with N1KO (NG, 1.27 +/-0.06; N1KO, 1.26 +/-0.04; p=0.8112). **K)** The proportion of aligned PSD-95 NCs is unaltered in N1KO synapses (NG, 18.91 +/-3.12%; N1KO, 21.79 +/-2.18%; p=0.1797). All data mean +/-SEM.

Beyond the properties of individual NCs, the average number of PSD-95 NCs per synapse was also increased in GluN1 knockout neurons (Fig 2F). As PSD-95 NC number scales with PSD size (Dharmasri et al., 2023; Broadhead et al., 2016), we wondered if this was simply a consequence of the enlarged PSD in GluN1 knockout synapses. A linear regression analysis indicated that while the slope in the relationship between PSD size and PSD-95 NC number was unchanged, GluN1 knockout PSDs contained more PSD-95 NCs regardless of synapse size (Fig 2G). Ultimately, despite the increase in PSD-95 NC number, the reduced NC volume and the enlarged PSD result in a smaller proportion of the PSD occupied by PSD-95 NCs (Fig 2H).

Since an important property of PSD-95 NCs is their trans-synaptic alignment with release sites (Tang et al., 2016), we wondered if this reduction in the proportion of the PSD that contains NCs would weaken the likelihood of nanocolumn alignment. We tested this with two measures of alignment. First, we measured the peak-to-peak distance between PSD-95 NCs and their closest Munc13-1 NC and found that there was no difference based on genotype (Fig 2I). We next measured the normalized density of Munc13-1 across from PSD-95 NCs as an indicator of trans-synaptic enrichment and observed no difference between control and GluN1 knockout neurons (Fig 2J). Consistent with this, the proportion of PSD-95 NCs that were aligned with Munc13-1 high density regions remained unchanged in GluN1 knockout synapses (Fig 2K). Together, this data suggests that PSD-95 NC trans-synaptic alignment is preserved in GluN1 knockout neurons.

### Postsynaptic Manipulation of NMDAR Content Restructures the Active Zone

Despite causing changes to PSD size and PSD-95 nanostructure, the loss of NMDARs did not disrupt alignment of PSD-95 NCs with Munc13-1. This, coupled with our confocal imaging observations, suggests that the AZ may have been reorganized similarly to the PSD in the GluN1 knockout synapses. Super-resolution imaging of Munc13-1 (Fig 3A) indeed revealed that the AZ, defined as the region bounded by Munc13-1 localizations, was enlarged (Fig 3B) by a similar factor (1.95x) to the PSD (1.90x). When assessing the nanoscale organization of Munc13-1, autocorrelation analysis revealed an increase in magnitude over short shift radii after GluN1 knockout (Fig 3C), predictive of a higher protein density within Munc13-1 NCs. This was further supported by a higher peak magnitude of the Munc13-1 autoenrichment (Fig 3D) and no change to Munc13-1 NC volume (Fig 3E). These data suggest that the organization of presynaptic Munc13-1 molecules into NCs is tuned by the presence of postsynaptic NMDARs.

Consistent with this idea, we observed an increase in the average number of Munc13-1 NCs per AZ in GluN1 knockout neurons (Fig 3F). As the number of PSD-95 NCs was increased regardless of PSD size in GluN1 knockout synapses, we tested if a similar effect was observed in the active zone. However, linear regression revealed a different effect – NMDAR loss yielded a shallower relationship between AZ size and Munc13-1 NC number, such that smaller synapses in GluN1 knockout neurons contained more Munc13-1 NCs than the control condition, while larger synapses contained fewer Munc13-1 NCs (Fig 3G). This change, coupled with most AZs being enlarged in the GluN1 knockout synapses, resulted in a stark reduction in the average proportion of the AZ occupied by Munc13-1 NCs (Fig 3H).

**Figure 3.**
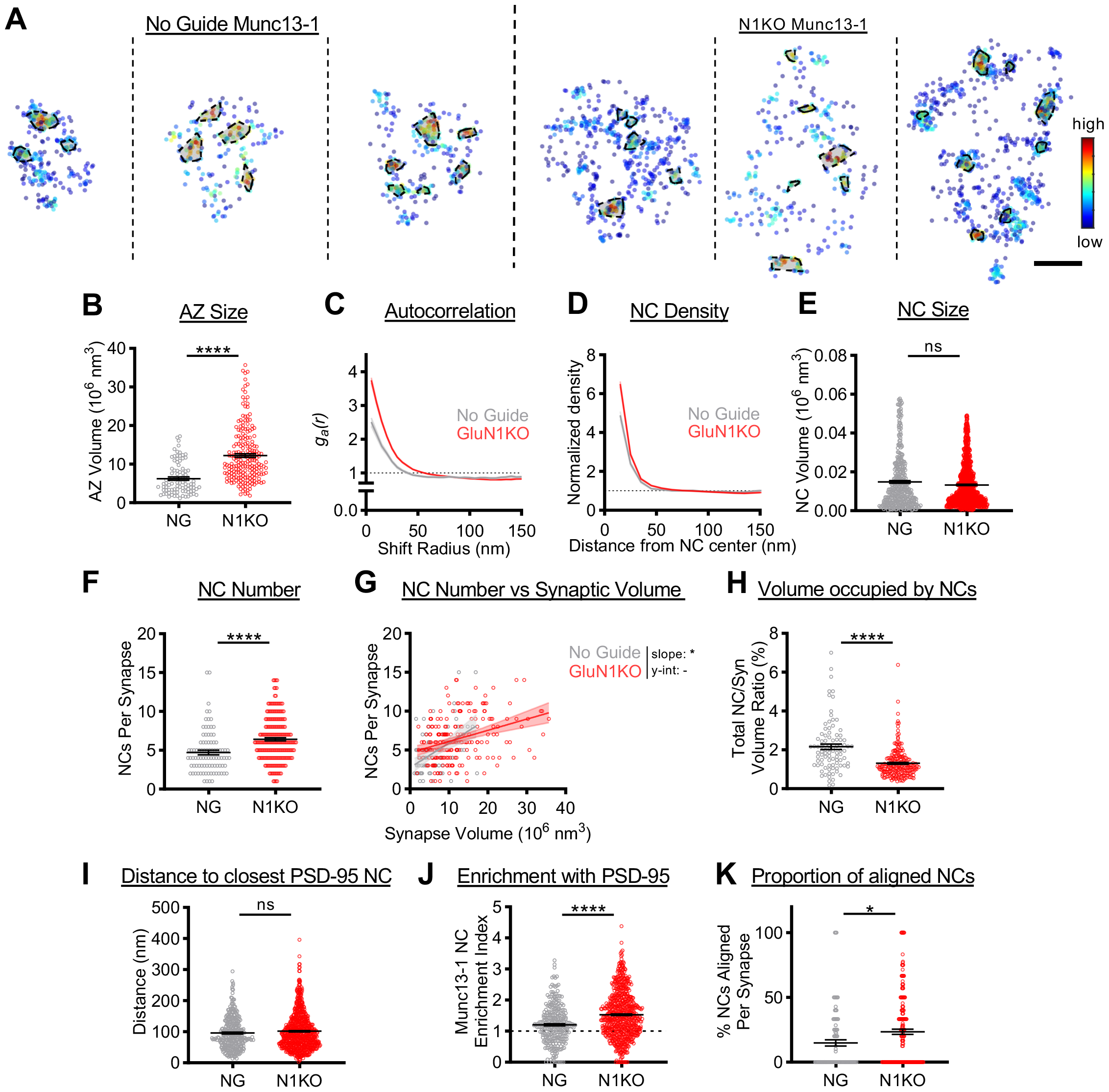
Trans-synaptic alteration of Munc13-1 organization in GluN1 knockout synapses. **A)** Examples of 3D DNA-PAINT Munc13-1 synaptic clusters from (Left) NG and (Right) N1KO synapses. Peak density-normalized local density indicated by heat maps. Detected NCs indicated by shaded shapes. Scale bar: 100 nm. Data in B,C,F-H are all synapses (NG, n = 91 synapses; N1KO, n = 213 synapses). Data in D,E,I-K are all NCs (NG, n = 372 NCs; N1KO, n = 1202 NCs). **B)** Munc13-1 synaptic clusters are enlarged at N1KO synapses (NG, 6.27 +/-0.42 10^6^ nm^3^; N1KO, 12.25 +/-0.48 10^6^ nm^3^; p<0.0001). **C)** Autocorrelation predicts denser Munc13-1 NCs in N1KO synapses. **D)** Average Munc13-1 NC has increased peak density in N1KO synapses. **E)** N1KO does not alter Munc13-1 NC volume (NG, 0.015 +/-0.00073 10^6^ nm^3^; N1KO, 0.013 +/-0.00032 10^6^ nm^3^; p=0.6395). **F)** N1KO synapses contain more Munc13-1 NCs (NG, 4.71 +/-0.28 NCs/synapse; N1KO, 6.43 +/-0.20 NCs/synapse; p<0.0001). **G)** N1KO alters relationship between AZ size and Munc13-1 NC number (NG, slope: 0.31, y-intercept: 2.75, R^2^: 0.22; N1KO, slope: 0.14, y-intercept: 4.67, R^2^: 0.12. slope p=0.0204, y-intercept cannot be tested due to the change in slope). **H)** Less synaptic volume occupied by Munc13-1 NCs in N1KO synapses (NG, 2.16 +/-0.14% volume; N1KO, 1.31 +/-0.05% volume; p<0.0001). **I)** Munc13-1 NC distance to closest PSD-95 NC is unchanged with N1KO (NG, 96.08 +/-2.82 nm; N1KO, 101.8 +/-2.04 nm; p=0.3083). **J)** Munc13-1 NCs are more enriched with PSD-95 in N1KO synapses (NG, 1.21 +/-0.04; N1KO, 1.53 +/-0.03; p<0.0001). **K)** N1KO increases the proportion of aligned Munc13-1 NCs (NG, 14.90 +/-2.45%; N1KO, 23.49 +/-1.93%; p=0.0123). All data mean +/-SEM.

Given these organizational changes to the AZ, we considered whether Munc13-1 NCs were organized differently with respect to their postsynaptic counterparts. The peak-to-peak distance between Munc13-1 NCs and their closest PSD-95 NCs was not different between conditions (Fig 3I), in agreement with the PSD-95 data that alignment was preserved. When we measured the normalized density of PSD-95 across from Munc13-1 NCs, we observed an increased enrichment in GluN1 knockout synapses (Fig 3J), consistent with preserved alignment and increased PSD-95 density around NCs (Fig 2C-D). Intriguingly, the loss of NMDARs increased the proportion of Munc13-1 NCs that were aligned with PSD-95 (Fig 3K). These data suggest that not only is Munc13-1 NC alignment preserved after GluN1 knockout, but the loss of NMDARs results in more robust organization of release sites across from regions of high local PSD-95 density within the synapse, likely to strengthen synaptic transmission. Altogether, these data demonstrate that NMDAR loss drives retrograde, trans-synaptic reorganization of presynaptic nanostructure.

### Prolonged NMDAR Antagonism Does Not Fully Recapitulate Munc13-1 Reorganization

Genetic deletion of GluN1 eliminates not only the structural role receptors may play in the synapse, but also the important functions mediated by NMDAR activation. To disambiguate these roles, we pharmacologically blocked NMDAR activity in wild type cultures. In our hands, the LCv2 GluN1 knockout reached completion between the second and third week post infection (Fig 1C), meaning that a proportion of the analyzed synapses analyzed may have only been without NMDARs for approximately one week. We therefore incubated our cultures in either 100 μM D,L-APV or vehicle from DIV 14 to DIV 21, refreshed every 48 hours prior to fixation, and tested for any changes to synaptic structure.

We observed with confocal microscopy that prolonged NMDAR antagonism induced overlapping, but distinct, changes compared to NMDAR loss (Fig 4A-B). APV treatment increased PSD-95 content at the synapse that was driven by an increase in PSD-95 density without a change to PSD size (Fig 4C). This is in contrast to the GluN1 knockout synapses, where the PSD increased in size with no change to protein density. Munc13-1 changes however were similar to those with GluN1 knockout, as Munc13-1 content increased due to an increase in AZ size and not a change in Munc13-1 density (Fig 4D, as compared to Fig 1G). Although the effect size was smaller than in GluN1 knockout synapses, this does suggest that the change we observed in AZ size with GluN1 loss can in part be attributed to the loss of NMDAR activity.

**Figure 4.**
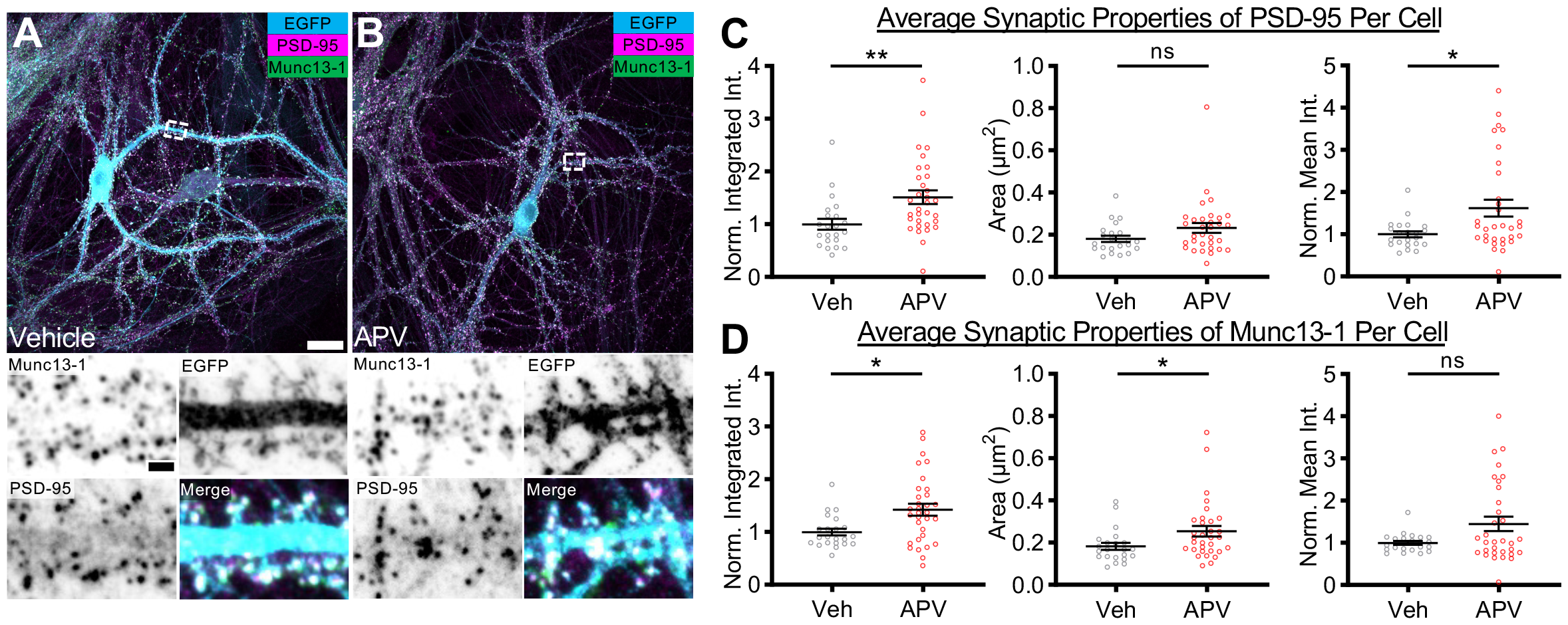
Prolonged NMDAR antagonism yields overlapping and distinct changes to synaptic properties compared to NMDAR loss. **A**,**B)** (Top) Example neurons in dissociated hippocampal cultures treated with either (A) vehicle (Veh) or (B) 100 μM D,L-APV (APV). Scale bar: 20 μm. (Bottom) Dendrite stretches from example cells above showing Munc13-1 staining, PSD-95 staining, cell identifying marker, and merged image. Scale bar: 2 μm. For C and D, per cell analyses were conducted to account for treatment (Veh, n = 22 cells; APV, n = 32 cells). **C)** Quantification of synaptic PSD-95 properties per cell. (Left) Normalized integrated intensity (Veh, 1.00 +/-0.10; APV, 1.51 +/-0.13; p=0.0012). (Middle) Puncta area (Veh, 0.18 +/-0.02 μm^2^; APV, 0.23 +/-0.02 μm^2^; p=0.0992). (Right) Normalized mean intensity (Veh, 1.00 +/-0.07; APV, 1.62 +/-0.20; p=0.0316). **D)** Quantification of synaptic Munc13-1 properties per cell. (Left) Normalized integrated intensity (Veh, 1.00 +/-0.07; APV, 1.43 +/-0.11; p=0.0125). (Middle) Puncta area (Veh, 0.18 +/-0.02 μm^2^; APV, 0.25+/-0.02 μm^2^; p=0.0162). (Right) Normalized mean intensity (Veh, 1.00 +/-0.05; APV, 1.45 +/-0.17; p=0.5124). All data mean +/-SEM.

Since the pharmacological manipulation resulted in similar changes to the AZ as in the knockout, we further investigated whether or which properties of Munc13-1 nano-organization were impacted by sustained antagonism of NMDAR activity. Autocorrelation analysis indicated that Munc13-1 was organized into denser NCs (Fig 5B). Consistent with this, the peak of the autoenrichment relationship was also increased (Fig 5C). Thus, presynaptic Munc13-1 NC properties are sensitive to NMDAR activity. The nature and magnitude of these effects were very similar to those following GluN1 knockout, further suggesting that they may be mediated by receptor activity, rather than receptor structure or protein interactions.

However, several other changes were observed following APV treatment that did not recapitulate the effects of GluN1 knockout. The steeper slope of the autocorrelation further suggested that Munc13-1 NCs may be smaller in APV-treated synapses. Indeed, detected Munc13-1 nanoclusters had a smaller volume following APV treatment (Fig 5D), unlike in knockout synapses. Interestingly, our super-resolution analysis indicated that prolonged APV treatment did not enlarge the AZ (Fig 5E). This is consistent with our confocal analysis showing the small effect size and higher cell-to-cell variability in synapse size in the APV-treated group (Veh CV: 43.45%; APV CV: 55.21%), suggesting that the lack of NMDAR activity may not completely account for the differences observed with total NMDAR loss. Also, in contrast to the GluN1 knockout synapses, prolonged APV treatment did not increase the number of Munc13-1 NCs (Fig 5F) or change the relationship between AZ size and Munc13-1 NC number (Fig 5G). While APV treatment did reduce the proportion of the AZ occupied by Munc13-1 NCs (Fig 5H), this appeared due to the smaller NC size, unlike in the knockout synapses. Thus, presynaptic organization appears sensitive to both the physical presence of NMDARs and to NMDAR-mediated signaling.

**Figure 5.**
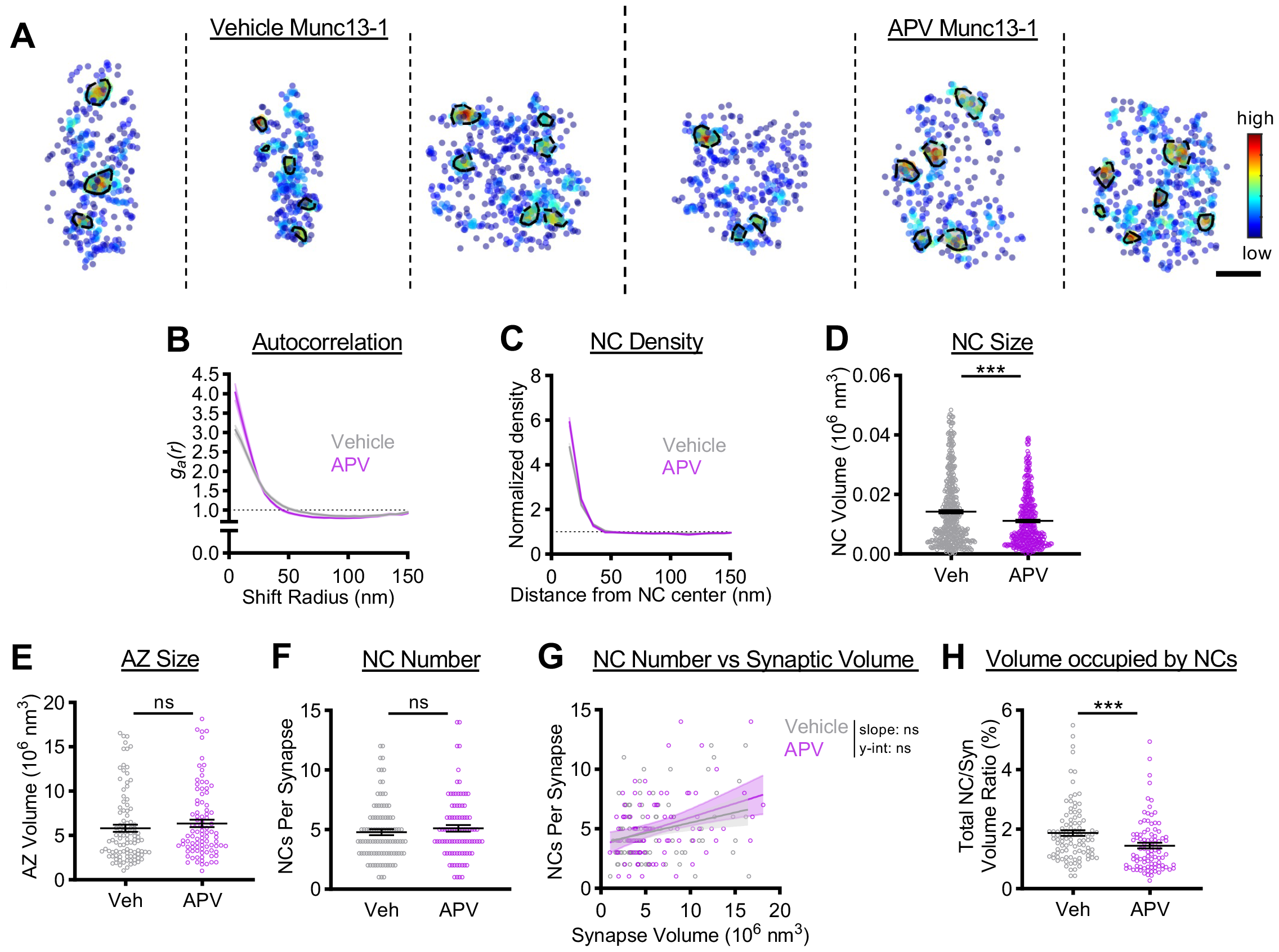
Loss of NMDAR activity does not fully recapitulate GluN1 knockout. **A)** Examples of 3D DNA-PAINT Munc13-1 synaptic clusters from (Left) Veh and (Right) APV treated synapses. Peak density-normalized local density indicated by heat maps. Detected NCs indicated by shaded shapes. Scale bar: 100 nm. Data in B,C,F-H are all synapses (Veh, n = 101 synapses; APV, n = 92 synapses). Data in D and E are NCs (Veh, n = 446 NCs; APV, n = 422 NCs). **B)** Autocorrelation predicts denser, smaller Munc13-1 NCs with APV treatment. **C)** APV treatment increases average Munc13-1 NC peak density. **D)** Munc13-1 NC volume is reduced with prolonged NMDAR silencing (Veh, 0.014 +/-0.00056 10^6^ nm^3^; APV, 0.011 +/-0.00043 10^6^ nm^3^; p=0.0007). **E)** Munc13-1 synaptic cluster size is unchanged with APV treatment (Veh, 5.81 +/-0.39 10^6^ nm^3^; APV, 6.37 +/-0.41 10^6^ nm^3^; p=0.1739). **F)** APV treatment does not impact Munc13-1 NC number per synapse (Veh, 4.79 +/-0.24 NCs/synapse; APV, 5.11 +/-0.28 NCs/synapse; p=0.3404). **G)** The relationship between AZ size and Munc13-1 NC number is unaltered by APV (Veh, slope: 0.17, y-intercept: 3.79, R^2^: 0.08; APV, slope: 0.23, y-intercept: 3.62, R^2^: 0.12. slope p=0.4850, y-intercept p=0.5606). **H)** Less synaptic volume occupied by Munc13-1 NCs with APV treatment (Veh, 1.87 +/-0.10% volume; APV, 1.45 +/-0.09% volume; p=0.0002).

## Discussion

In this study, we used a combination of confocal and super-resolution imaging to test the impact of NMDAR loss on synaptic structure. While the role of NMDARs as structural hubs for critical effector molecules led us to predict that their loss would result in less organized nanostructure, we instead observed the opposite. Genetic deletion of GluN1 resulted in a larger PSD, with PSD-95 forming denser, more numerous NCs than at control synapses. Surprisingly, loss of NMDARs had a retrograde effect on AZ organization, resulting in larger AZs with denser, more numerous NCs and a greater proportion of release sites aligned with PSD-95 NCs. These changes likely lead to stronger synaptic transmission in these synapses (Li et al., 2022). Notably, while prolonged antagonism of NMDAR activity yielded clear trans-synaptic effects on the density of Munc13-1 within NCs, it did not fully recapitulate the changes observed in GluN1 knockout synapses. Together, our data suggest that both NMDAR activity and structure play unique roles in maintaining both pre- and postsynaptic nano-organization.

Deletion of GluN1 yielded larger PSDs with smaller but denser and more numerous PSD-95 NCs. An enlarged PSD has been previously reported in GluN1 knockout synapses (Ultanir et al., 2007). In addition, GluN1 knockout results in stronger AMPAR-dependent EPSCs (Adesnik et al., 2008; Incontro et al., 2014), as well as increases in either mEPSC amplitude (Ultanir et al., 2007) or frequency (Adesnik et al., 2008; Ultanir et al., 2007). The smaller, denser PSD-95 NCs could function to concentrate receptors more tightly under release sites, yielding greater activation of AMPARs per release event (Freche et al., 2011; Savtchenko and Rusakov, 2014). Indeed, it is easily conceivable that the increase in PSD-95 NC number allows for more instances of receptor alignment to release sites within a synapse (Lisman and Raghavachari, 2006). Thus, the results of our structural study indicate changes on the level of the individual synapse that are consistent with previous reports and could potentially yield both stronger mEPSC amplitudes and an increase in mEPSC frequency.

Hippocampal neurons express predominantly GluN2A- and GluN2B-containing NMDAR subtypes. NMDAR subtypes may differentially regulate synaptic nanostructure, as they have different developmental expression profiles and protein interactions (Sanz-Clemente et al., 2013) and are present with distinct subsynaptic organizations (Anderson et al., 2023; Kellermayer et al., 2018). GluN1 knockout precludes our ability to distinguish how NMDAR subtype may be influencing the observed changes to PSD nanostructure. However, GluN2B knockouts display several characteristics of GluN1 knockouts, including stronger AMPAR-mediated EPSCs and increases in either mEPSC amplitude (Hall et al., 2007) or frequency (Gray et al., 2011). Furthermore, cultured hippocampal neurons largely express GluN2B-containing NMDARs (Metzbower et al., 2019; Sinnen et al., 2016), further suggesting that our observed changes in nanostructure may be due to the loss of GluN2B-containing NMDARs. Nevertheless, knockout of GluN2A has also been demonstrated to increase AMPAR-mediated mEPSC amplitude (Gray et al., 2011), and thus our observations may stem from separate effects due to the loss of each NMDAR subtype. Dissecting the role NMDAR subtypes may have in shaping synaptic nanostructure could expand our understanding of mechanisms of synaptic development and maturation.

Loss of NMDARs resulted in a retrograde reorganization of the AZ. The increase in Munc13-1 NC number suggests a larger readily releasable pool is present in GluN1 knockout synapses (Karlocai et al., 2021; Sakamoto et al., 2018). In addition, the increase in Munc13-1 density within NCs could suggest that docked vesicles at these synapses are more likely primed and ready for release (Aldahabi et al., 2022). This, in addition to the increased proportion of release sites that are aligned with PSD-95 NCs in the GluN1 knockout synapses, suggests synaptic transmission is enhanced in synapses lacking NMDARs. Thus, one potential conclusion from these observations is that in wildtype synapses, the presence of NMDARs may act to limit the strength of individual AZs.

The results of our APV treatment experiments further support that NMDARs suppress AZ strength. Prolonged antagonism of NMDAR activity increased Munc13-1 NC density compared to vehicle treatment, demonstrating that NMDAR activity does influence AZ organization. This increase came with a decrease in NC volume, suggesting that loss of NMDAR activity may have resulted in a “contracted” Munc13-1 NC. This deviates from GluN1 knockout synapses, which showed an increased density and no change to Munc13-1 NC volume and which may be indicative of more Munc13-1 packed into NCs in the total absence of the receptor. Together, this suggests that the confluence of NMDAR activity and presence at the synapse may be important to suppress vesicle priming in the AZ by regulating characteristics of Munc13-1 NCs.

One intriguing unknown is the molecular route by which the NMDAR complex mediates presynaptic nano-organization. Previous work has focused on how certain molecular interactions and pathways are important for promoting NMDAR retention and activity at the synapse. For instance, extracellular interactors such as EphB2 (Dalva et al., 2000; Hanamura et al., 2017; Washburn et al., 2020), Nlgn1 (Budreck et al., 2013), and SALM1 (Wang et al., 2006) can impact synaptic NMDAR levels. In addition, NMDAR activity is indirectly regulated by presynaptic cell adhesion molecules, such as Nrxn1 (Dai et al., 2021, 2019) and LAR-RPTPs (Kim et al., 2020; Lie et al., 2021, 2016; Sclip and Südhof, 2020). While these interactors and pathways influence synaptic NMDAR activity, it is untested whether any of these mechanisms are ‘bidirectional’ – in other words, can NMDARs signal through the same molecular networks that are responsible for recruiting them? If so, they may represent potential avenues by which the NMDAR complex can modulate Munc13-1 nanostructure and thereby regulate AZ function.

It is important to note that previous studies of GluN1 deletion have shown no changes to the mean presynaptic probability of release assessed physiologically (Adesnik et al., 2008; Gray et al., 2011; Incontro et al., 2014). Previous modeling has suggested that probability of release is influenced by the size of the readily releasable pool (Sun et al., 2005), and thus the increase in the average number of Munc13-1 NCs we observed would be expected to produce an increase in release probability. However, we found in the GluN1 knockout synapses that despite the increase in the average NC number, smaller AZs had more Munc13-1 NCs and larger AZs had fewer Munc13-1 NCs, when compared to control (Fig 3G). Thus, our data predicts a heterogeneous impact of NMDAR loss on release probability on a per synapse basis, where larger synapses may have a lower release probability than smaller synapses. Indeed, imaging experiments that can measure activity at individual synapses document that spontaneous release rate and evoked release probability vary widely across hippocampal synapses (Jensen et al., 2021; Metzbower et al., 2019), while this heterogeneity is lost in typical physiological recordings which average the behavior of many synapses. Revisiting the impact of NMDAR loss on presynaptic release probability with optical measurements capable of single synapse resolution may be able to detect previously unappreciated roles of NMDARs on presynaptic function.

The NMDAR complex is a key player in synaptic dynamics (Lüscher and Malenka, 2012). Notably, NMDAR-dependent synaptic plasticity is associated with changes to synaptic nanostructure. NMDAR-dependent LTP results in changes to PSD-95 organization (Hruska et al., 2022, 2018), including PSD-95 NC enlargement (Clavet-Fournier et al., 2023) and an increase in NC density and alignment with release sites (Tang et al., 2016). NMDAR-driven LTD results in a reduction of PSD-95 NC number and release site alignment (Tang et al., 2016). NMDAR activity also impacts presynaptic nanostructure, as treatment with NMDAR antagonist APV for 48 hours decreases presynaptic clustering (Glebov et al., 2017). This ability of nanostructure to shift in accordance with NMDAR activity suggests a spectrum of synaptic nano-organization over which NMDARs hold considerable sway. From this perspective, the observed nanostructure of GluN1 knockout synapses may represent an extreme of such a spectrum, where the synapse is highly organized and aligned. We thus speculate that the NMDAR complex may play a role in controlling synaptic transmission via maintaining the dynamic range of synaptic nanostructure.

The results of this study may also shed light on synaptic rearrangement that occurs during pathological states associated with a decrease either in NMDAR number or activity at the synapse. Several examples of pathological NMDAR downregulation include: NMDAR hypofunction in schizophrenia (Balu, 2016), NMDAR antibody binding induced receptor internalization in Anti-NMDAR encephalitis (Dalmau and Graus, 2018), as well as reduction of NMDAR activation (Sinnen et al., 2016) and increased endocytosis (Kurup et al., 2010) in Alzheimer’s disease models involving Amyloid β, which was recently and independently associated with synaptic nanostructural rearrangement (Zhu et al., 2023). Our results suggest that disease states that differentially impact NMDAR function or presence at the synapse may yield distinct nanostructural states that contribute to the unique functional status of synapses in these disorders. This is further highlighted by the ability of the NMDAR complex to host multiple modes of signaling (Dore et al., 2017; Park et al., 2022). Whether it is through contributing to postsynaptic current, enabling activity-dependent synaptic calcium flux, docking of critical effectors such as CaMKII or non-ionotropic signaling proteins, or binding cleft-resident molecules, there are several potential avenues through which NMDARs can impact synaptic nanostructure. We suggest that the disruption of specific combinations of such processes underpins the diversity in NMDAR-centric pathologies. Understanding the precise mechanisms by which NMDARs influence subsynaptic protein organization represents a potentially powerful lens through which to study both synaptic transmission and synaptic disruption in various diseases and disorders.

## Acknowledgments

The authors would like to thank the members of the Blanpied Lab for helpful discussion and evaluation of the manuscript, as well as Minerva Contreras for significant technical support.

